# A Genetic Pathway Composed of *EDT1/HDG11*, *ERECTA*, and *E2Fa* Loci Regulates Water Use Efficiency by Modulating Stomatal Density

**DOI:** 10.1101/232801

**Authors:** Xiao-Yu Guo, Yao Wang, Ping Xu, Guo-Hua Yu, Li-Yong Zhang, Yan Xiong, Cheng-Bin Xiang

## Abstract

Improvement of crop drought resistance and water use efficiency (WUE) has been a major endeavor in agriculture. ERECTA is the first identified major effector of water use efficiency. However, the underlying molecular mechanism is not well understood. Here, we report a genetic pathway, composed of *EDT1/HDG11*, *ERECTA*, and *E2Fa* loci, which regulates water use efficiency by modulating stomatal density. The HD-START transcription factor EDT1/HDG11 transcriptionally activates *ERECTA* expression by binding to an HD *cis*-element in the *ERECTA* promoter. ERECTA in turn relies on E2Fa to control the expression of cell-cycle related genes and the transition from mitosis to endocycle, which leads to increased nuclear DNA content in leaf cells, and therefore increased cell size and decreased stomatal density. The decreased stomatal density improves plant WUE. Our study demonstrates the EDT1/HDG11-ERECTA-E2Fa genetic pathway that reduces stomatal density by increasing cell size, providing a new avenue to improve WUE of crops.

## Introduction

Drought stress is one of the most severe environmental constraints that greatly restrict plant growth, distribution and crop production (Zhu, 2002; Kissoudis et al., 2016). An effective strategy of drought resistance used by plants is reducing transpirational water loss, which allows plants to maintain an adequate water status to sustain critical physiological and biochemical processes (Chaves et al., 2003). However, a reduction in transpirational water loss often leads to a decline in biomass accumulation because of reduced carbon assimilation. Therefore, plant water use efficiency (WUE), which is defined as plant production per amount of water used, is critical to plant survival and crop yield (Yoo et al., 2009).

*ERECTA*, the first reported WUE-related gene, was isolated by quantitative trait locus (QTL) analysis for carbon isotope discrimination (Masle et al., 2005). Altering the *ERECTA* gene expression displayed significant change of plant WUE (Masle et al., 2005; Shen et al., 2015). However, the causal molecular mechanism remains unclear. *ERECTA* encodes a leucine-rich repeat receptor-like kinase (LRR-RLK) (Torii et al., 1996) with many important functions in plant development (Xu et al., 2003; Shpak et al., 2004; Douglas and Riggs, 2005; Meng et al., 2012; Pillitteri and Torii, 2012; Bemis et al., 2013; Cui et al., 2014). ERECTA and its two highly homologous receptor-like kinases (RLKs), ER-LIKE1 (ERL1) and ER-LIKE2 (ERL2), synergistically inhibit stomatal differentiation and enforce proper stomatal spacing (Shpak et al., 2005; Lau and Bergmann, 2012; Lee et al., 2012; Lee et al., 2015; Meng et al., 2015; Qi et al., 2017).

Unlike stomatal patterning defective mutants, the *ERECTA* single-gene knockout mutant *er-105* exhibits increased stomatal density. When overexpressed, *ERECTA* reduced stomatal density and increased WUE without changing stomatal patterning (Masle et al., 2005; Shpak et al., 2005).Similar phenotype is exhibited by the *edt1* mutant we previously reported (Yu et al., 2008). In *edt1* mutant, the homeodomain leucine zipper (HD-Zip) transcription factor *EDT1/HDG11* is constitutively activated by an activation tagging T-DNA insertion, leading to reduced stomatal density and improved WUE but unaltered stomatal index (number of stomata per total number of epidermal cells) (Yu et al., 2008). In the wild type, *EDT1/HDG11* is weakly expressed in reproductive organs. Knockout of *EDT1/HDG11* did not produce any significantly different phenotype from the wild type (Yu et al., 2008). However when overexpressed, *AtEDT1/HGD11* reduced stomatal density and improved WUE in Arabidopsis and tobacco (Yu et al., 2008), and rice (Yu et al., 2013). *AtEDT1/HGD11*-conferred drought resistance and improved WUE appear to be well conserved in higher plants since the *edt1* mutant phenotypes could be recapitulated by overexpressing *AtEDT1/HDG11* in cotton and poplar (Yu et al., 2016), wheat (Li et al., 2016), turf grass (Cao et al., 2009), sweet potato (Ruan et al., 2012), pepper (Zhu et al., 2015), Chinese kale (Zhu et al., 2016), *Salvia miltiorrhiza* (Liu et al., 2017), and alfalfa (Zheng et al., in press).

The Arabidopsis epidermal patterning factor (EPF) (Kondo et al., 2010; Lee et al., 2015) mutants with altered stomatal density were elegantly used to test the relationship between WUE and stomatal density. It was found that reduced stomatal density was correlated with increased WUE (Doheny-Adams et al., 2012). Furthermore, stomatal density can be genetically manipulated to improved WUE (Franks et al., 2015; Hepworth et al., 2015).

Transpiration and CO_2_ uptake are the two determinants of WUE. Both occur primarily through stomata and are modulated by stomatal movements and stomatal density (Hetherington and Woodward, 2003; Yoo et al., 2010). The WUE in *er-105* and *edt1* is altered because of altered stomatal density, which is caused by the change of epidermal cell size (Masle et al., 2005; Yu et al., 2008).

The final cell size within an organ is often, but not always, correlated with the ploidy level resulting from the nuclear endoreduplication (Breuer et al., 2010). Transcription factor E2Fa is a key regulator in cell cycle-endocycle progression. Ectopic expression of *E2Fa* has an opposite effect on cell proliferation during *Arabidopsis* development, which increases cell number in the cotyledons (De Veylder et al., 2002) while it decreases cell number in mature leaves (He et al., 2004). Further study indicated that E2Fa has a dual regulatory role. Overexpression of *E2Fa* in *Arabidopsis* can stimulate cell proliferation via activating S-phase genes expression and inducing extra round of DNA replication (endoreduplication). Through binding or dissociating with retinoblastoma-related protein (RBR), E2Fa can control cell cycle and endocycle processes (Magyar et al., 2012).

In this report, we continue to investigate *AtEDt1/HDG11*-conformed drought resistance and improved WUE and provide evidences to suggest a genetic pathway composed of EDT1/HDG11, ERECTA, and E2Fa as key components in regulating stomatal density and WUE. *ERECTA* is a major target of EDT1/HDG11 and is transcriptionally upregulated by the EDT1/HDG11, which consequently enhances E2Fa activity leading to a higher polyploidy level in leaf cells. The elevated ploidy levels increase leaf cell size and thus decrease stomatal density, which consequently improves plant WUE.

## Results

### *EDT1/HDG11* depends on *ERECTA* in regulating stomatal density and WUE

To determine the genetic relationship between *ERECTA* and *EDT1/HDG11*, we introduced the activated *EDT1/HDG11* from a gain-of-function mutant *edt1* into *er-105* background by crossing and generated a double mutant *edt1er-105*, in which *EDT1/HDG11* is overexpressed while *ERECTA* is completely knocked out (χ2 = 5.86, 0. 05 ≤ P ≤ 0.1). The double mutant *edt1er-105* displayed similar phenotype to *er-105* (Fig. 1 and Supplemental Fig. 1). In the double mutant *edt1er-105*, leaf cell density and stomatal density were similar to the level of *er-105* (Fig. 1a and b). WUE was significantly decreased compared to that of *edt1* mutant (Fig. 1c), which was contributed by increased transpiration rate and slightly decreased photosynthesis rate (Supplemental Fig. 1c and d). These results suggest that transpiration rate and water use efficiency are correlated with changes of stomatal density, and that *EDT1/HDG11* functions through *ERECTA*, since knockout of *ERECTA* is sufficient to block the function of *EDT1/HDG11* in regulating stomatal density and water use efficiency.

**Figure 1.**
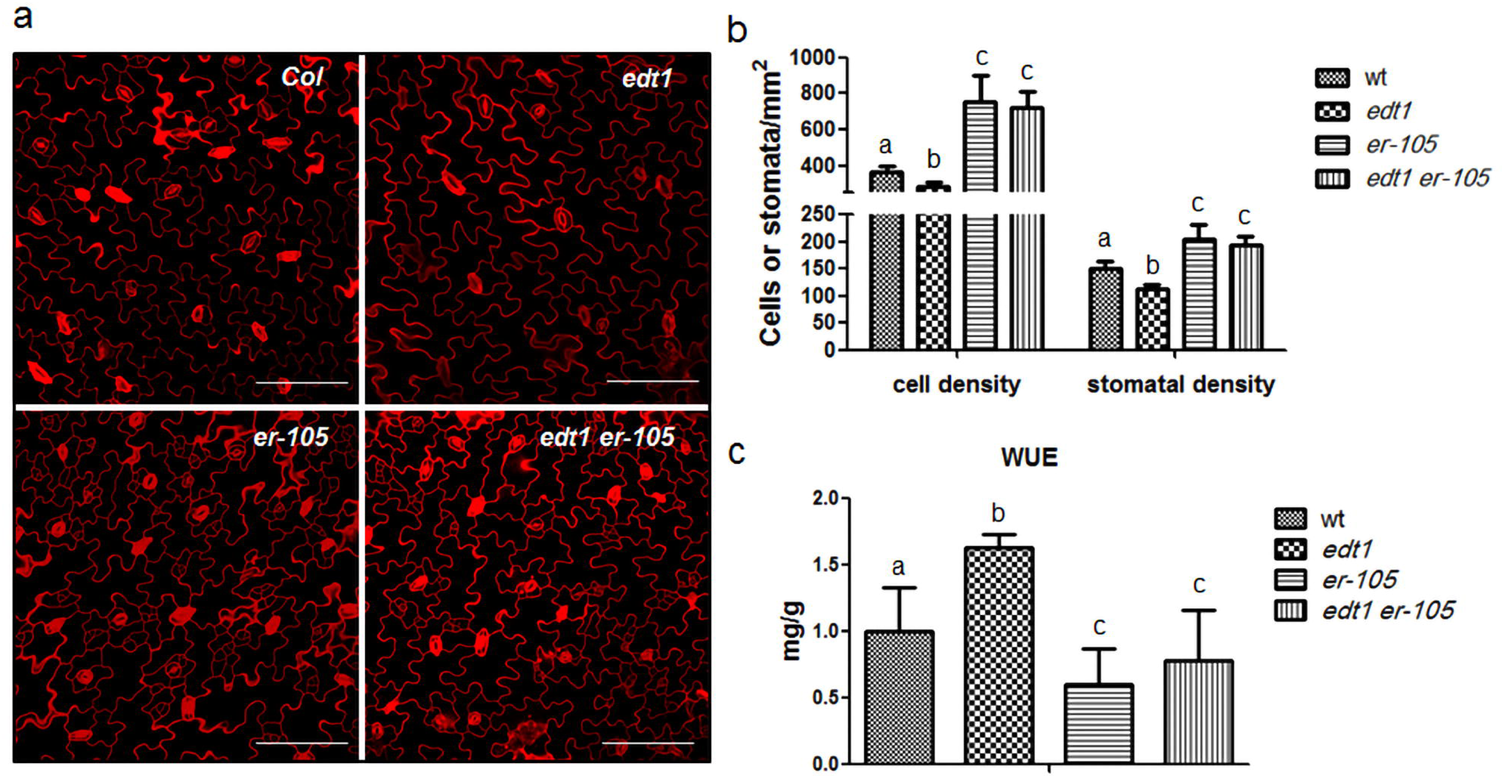
EDT1/HDG11 is dependent on ERECTA in regulating stomatal density and WUE. (a)Confocal images of adaxial cotyledon epidermis from 7-day-old wild type, *edt1*, *er-105*, and *edt1er-105*. Scale bar represents 100 μm. (b)Quantitative analysis of adaxial cotyledon cell density and stomatal density in 7-day-old wild type, *edt1, er-105*, and *edt1er-105*. Values represent the average (± SD) (n = 30), letters indicate differences statistically significant among the genotypes (p < 0.05; two-way ANOVA with Bonferroni post-test). (c)Comparison of WUE among the wild type, *edt1*, *er-105*, and *edt1er-105*. WUE was measured as described in Methods. Values are mean ± SD (n = 10 plants per genotype). Letters indicate statistical significance from one-way ANOVA followed by Bonferroni’s multiple comparison test (P < 0.05).

### Leaf cell size is positively correlated with polyploidy level

Stomatal density is determined largely by changes in epidermal cell size (Carins Murphy et al., 2016). To test whether the change of leaf cell size is accompanied by a change in leaf ploidy level via nuclear endoreduplication, we compared the DNA ploidy level of the fifth rosette leaf (which has the similar leaf size in different mutants) of four-weeks-old plants by flow cytometry. The ploidy distributions revealed significant differences between *edt1* with *er-105* and *edt1er-105*. The ploidy of *edt1* cells ranged from 2C to 32C with a peak at 8C, whereas the ploidy of the cells in *er-105* and *edt1er-105* were from 2C to 32C with a peak at 4C (Fig. 2a). The average percentage of high polyploidy (8C or higher) cells were 45.95%, 57.82%, 34.54% and 35.04% for the wild type, *edt1*, *er-105*and *edt1er-105*, respectively (Fig. 2b). The population of leaf cells with 8C and higher DNA ploidy level were significantly increased in edt1compared to *er-105* and *edt1er-105*. The cell size of the mutants positively correlated with their ploidy level, suggesting that an activated expression of *EDT1/HDG11* in *edt1* stimulates endoreduplication. The leaf ploidy level distribution of *edt1er-105* was similar to that of *er-105*, consistent with our hypothesis that the function of *HGD11* is dependent on *ERECTA* in regulating stomatal density and WUE. These results indicate that the enlarged cell size in leaves of the *edt1* mutant is correlated with the increase of ploidy levels, and *ERECTA* is required in this regulation.

**Figure 2.**
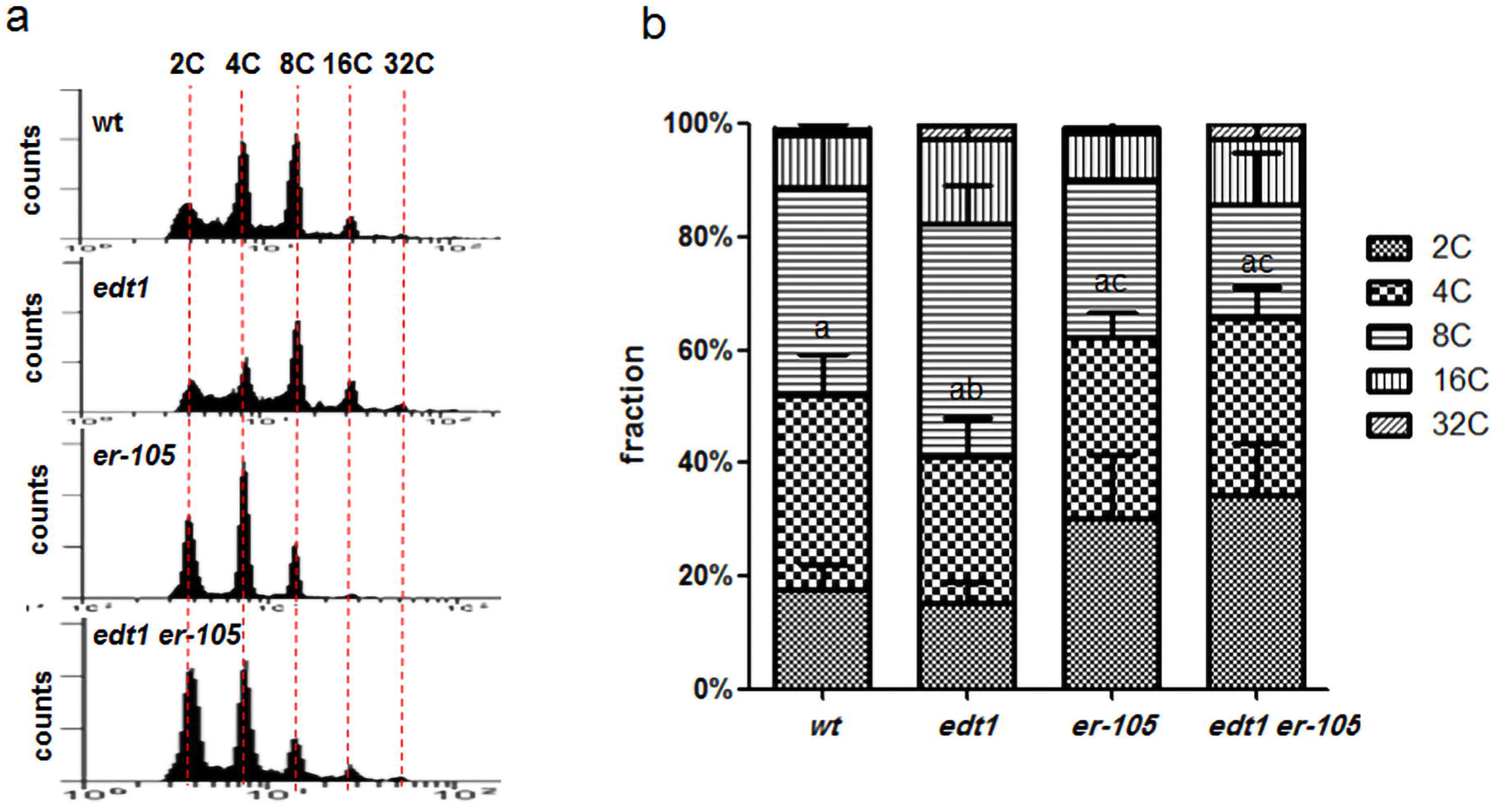
Leaf cell size is positively correlated with polyploidy level. (a)Ploidy histogram for nuclei isolated from the fifth rosette leaf of wild type, *edt1*, *er-105*, and *edt1er-105*. Note that the proportion of nuclei in the *edt1* peak at 8C, and peak at 4C in *er-105* and *edt1er-105*. Data are from one out of three independent experiments, which gave similar results. (b)Ploidy level distribution of wild type, *edt1*, *er-105* and *edt1er-105*. The nuclei isolated from the fifth rosette leaf of four-weeks-old plants were measured by flow cytometry. 10, 000 nuclei were counted for each sample. Data represent average ± SD (biological replicates of n=3; 6 leaves from at least 3 different plants per genotype). Significant differences are calculated by one-way ANOVA followed by Bonferroni’s multiple comparison test (P≤ 0.05).

### EDT1/HDG11 upregulates the expression of *ERECTA*

Rosette leaf transcriptome comparison between the wild type and *edt1* showed that the expression level of *ERECTA* was higher in *edt1* (Supplemental Fig. 2a). The higher expression level of *ERECTA* in *edt1* was also confirmed by quantitative realtime PCR (qRT-PCR) (Fig. 3a and Supplemental Fig. 2b). This result suggests that *ERECTA* might be transcriptionally upregulated by EDT1/HDG11 in *edt1* mutant.

**Figure 3.**
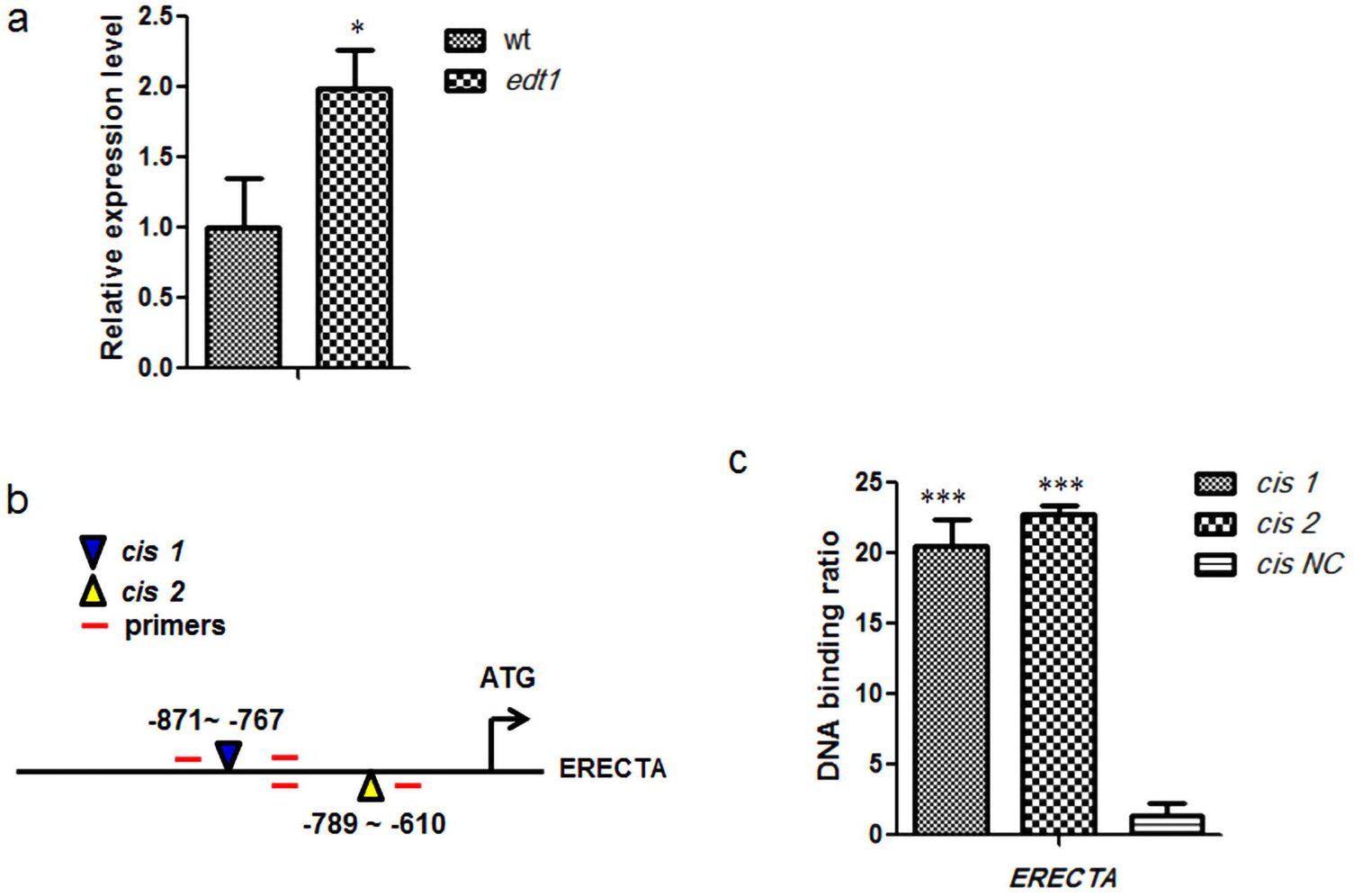
EDT1/HDG11 upregulates *ERECTA* expression by binding its promoter region. (a)Quantitative real-time PCR analysis of *ERECTA* expression in wild type and *edt1*. All assays were carried out at least three times and levels of statistical significance were calculated using student’s *t*-test (* P≤ 0.05). (b)The schematic illustration of the locations of *cis*-element 1 and *cis*-element 2 (inverted triangles) in the promoter of *ERECTA* and the primers used for quantitative RT–PCR (red short lines). (c)EDT1/HDG11 binds the promoter of *ERECTA in vivo*. ChIP-qPCR analysis was performed in the *35S⸬HA-HDG11* transgenic plants using HA tag-specific monoclonal antibody for immunoprecipitation. EDT1/HDG11 binds the specific *cis*-elements (*cis1* and *cis2*) on the *ERECTA* promoter but doesn’t bind the *cis-NC*. Values are mean ± SD. (*n*= 3 experiments, **P ≤ 0.01, ***P ≤ 0.001, Student’s *t*-test).

To investigate whether EDT1/HDG11 directly binds to the promoter of *ERECTA*, we analyzed *ERECTA* promoter sequence and found two putative binding sites of HD class transcription factors, *cis*-element 1 (*cis1:* AAATTAGT) and *cis*-element 2 (*cis 2:* TAATAATTA) (Fig. 3b and Supplemental Fig. 2c). Yeast-one-hybrid assay showed that EDT1/HDG11 was able to bind the *cis*-element 1 and *cis*-element 2 in yeast cells (Supplemental Fig. 2d). We further performed a chromatin immunoprecipitation assay followed by qPCR (ChIP-qPCR) analysis using the *EDT1/HDG11* overexpression line as described (Xu et al., 2014). The results from ChIP-qPCR showed that the *cis*-element 1 and *cis*-element 2 were significantly enriched in the immunoprecipitate. For the negative control, we did not detect any significant enrichment of control sequence NC within the *ERECTA* 3’ UTR (Fig. 3c and Supplemental Fig. 2e). These results demonstrate that EDT1/HDG11 specifically binds the promoter of ERECTA and potentially activates the transcription of *ERECTA*.

Rosette leaf transcriptome comparison and qRT-PCR analysis showed that the expression levels of *ER-LIKE1*(*ERL1*) and *ER-LIKE2*(*ERL2*), two closely related RLKs of *ERECTA*, were not significantly higher in *edt1* than that in wild type (Supplemental Fig. 2a, b). Although promoter scanning results indicated that a *cis*-element 1 (*cis1:* AAATTAGT) is presented in *ERL2* promoter, and a variant *cis*-element 1 (*cis1m:* AAATTATT) in *ERL1* promoter (Supplemental Fig. 2c). The ChIP-qPCR experiment results showed that these two regions were not significantly enriched with anti-HA antibody (Supplemental Fig. 2e), suggesting that *ERL1* and *ERL2* are not the targets of EDT1/HDG11.

### E2Fa is a downstream target of ERECTA

Since cell size is positively correlated with ploidy level in the mutants *edt1* and *er-105*, we explored the genetic relationship between *ERECTA* and key transcriptional factors involved in cell cycle regulation and found that *E2Fa* might be a downstream target of this pathway. We analyzed potential epistatic interaction by crossing the *ERECTA* overexpression line *(ER)* into the *E2Fa* knockout mutant (*e2fa*) background. The decreased leaf cell density and decreased stomatal density of *ER* were completely repressed in *ERe2fa*, indicating that ERECTA alters cell size and stomatal density through E2Fa (Fig. 4a and 4b). The ploidy level of leaf cells measured by flow cytometry further suggested that the observed variation in cell size is positively correlated with endoreduplication. Average percentage of high polyploidy cells with 8C and higher DNA ploidy level for the wild type and ER were 51.6% and 65.83% in the fifth rosette leaf, respectively, while it was 40.97% in *e2fa*. In contrast to *ER* plants, the high ploidy cell proportion of the double mutant *ERe2fa* was reduced to 40.58%, which was similar to that of *e2fa* (Fig. 4c). These results show that cell size is positively correlated with cell ploidy level as previously shown for *edt1* and *er-105* mutants (Fig. 2). Therefore, E2Fa is required for endoreduplication in *ER* plants. Moreover, plant growth morphology of the double mutant *ERe2fa* was similar to *e2fa* but not to *ER* (Supplemental Fig. 3). WUE of *ERe2fa* was similar to *e2fa* and lower than that of *ER* (Fig. 4d). These results indicate that E2Fa is a downstream target of ERECTA. ERECTA functions through E2Fa to control cell size, stomatal density, and WUE.

**Figure 4.**
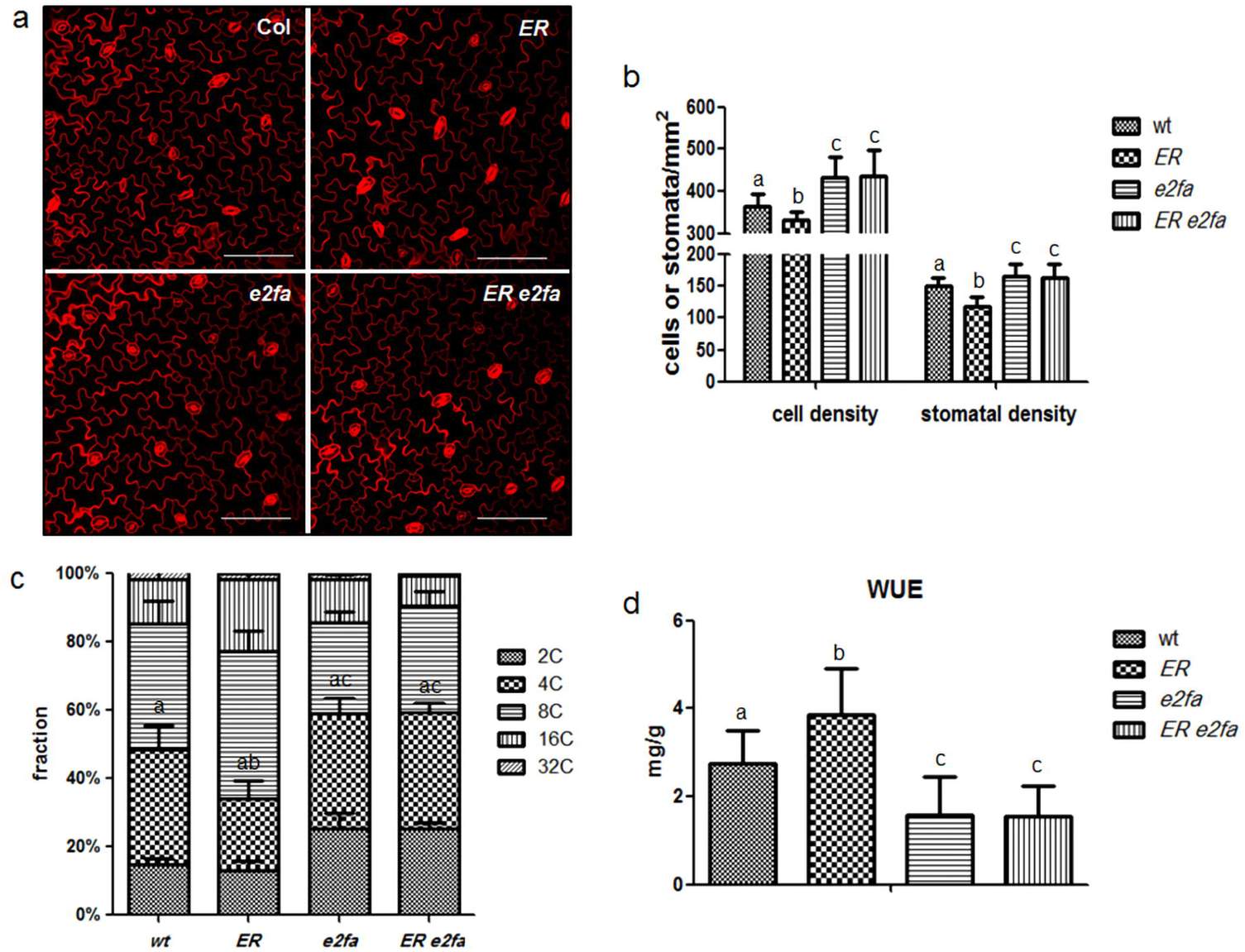
Genetic analyses show that ERECTA and E2Fa share a common pathway to regulate cell size, stomatal density and WUE. (a) Confocal images of adaxial cotyledon epidermis from 7-day-old wild type, *ER*,*e2fa* and *ERe2fa*. Scale bar represents 100 μm. (b) Quantitative analysis of adaxial cotyledon cell density and stomatal density in 7-day-old wild type, *ER*, *e2fa* and *ERe2fa*. Values are mean ± SD (n = 30), letters indicate differences statistically significant among the genotypes (p ≤ 0.05; two-way ANOVA with Bonferroni post-test). (c) Enlarged cell size is correlated with the increased polyploidy level in leaf cells. Ploidy level distribution of the fifth rosette leaf in four-week-old wild type, *ER*, *e2fa* and *ERe2fa* were measured by flow cytometry. 10, 000 nuclei were counted for each sample. Data represent average ± SD (biological replicates of n = 3; 6 leaves from at least 3 different plants per genotype). Significant differences are calculated by one way ANOVA followed by Bonferroni’fs multiple comparison test (p≤0.05). (d) Comparisons of WUE in wild type, *ER*, *e2fa* and *ERe2fa*. WUE was measured as described in Methods. Values are mean ± SD (n = 10 plants per genotype). Letters indicate statistical significance from one-way ANOVA followed by Bonferroni’s multiple comparison test (P ≤ 0.05).

### Expression levels of E2Fa target genes were changed in *ERECTA* mutants

To further confirm that ERECTA affects cell size through E2Fa, we measured the relative transcript levels of known E2Fa target genes that are required for G1/S transition and cell cycle progression by qRT-PCR in the wild type and *ERECTA* mutants. The target genes include *MINOCHROMOSOME MAINTENANCE 5*(*MCM5*) and *CELL DIVISION CONTROL 6* (*CDC6*), which are essential for the initiation of the DNA replication; *CYCLIN-DEPENDENTKINASEB1;1*(*CDKB1;1*) and *CYCLIN A2;3*(*CYCA2;3*), which play a central role in the control of mitotic cell cycle and inhibit the endocycle; *CELL CYCLE SWITCH PROTEIN 52 A1*(*CCS52A1*) and *CELL CYCLE SWITCH PROTEIN 52 A2*(*CCS52A2*) that play important regulatory roles in the transition from mitosis to endocycle by stimulating the degradation of mitotic cyclins. Compared to wild type, the expression of *CDKB1;1* and *CYCA2;3* were repressed in *ER*, but enhanced in *er-105*, *e2fa* and *ER e2fa*. The expression of endocycle activators *CCS52A* genes were increased in *ER*, but decreased in *er-105*,*e2fa* and *ERe2fa*. The expression of *MCM5* and *CDC6* were increased in *ER*, whereas decreased in *er-105*, *e2fa* and *ER e2fa* (Fig. 5). These results suggest that, through E2Fa transcription factor, ERECTA modifies the expression levels of cell cycle-related genes and regulates the transition from mitosis to endocycle, triggers cells to enter the endoreduplication cycle prematurely and eventually results in larger-sized cells with higher polyploidy levels.

**Figure 5.**
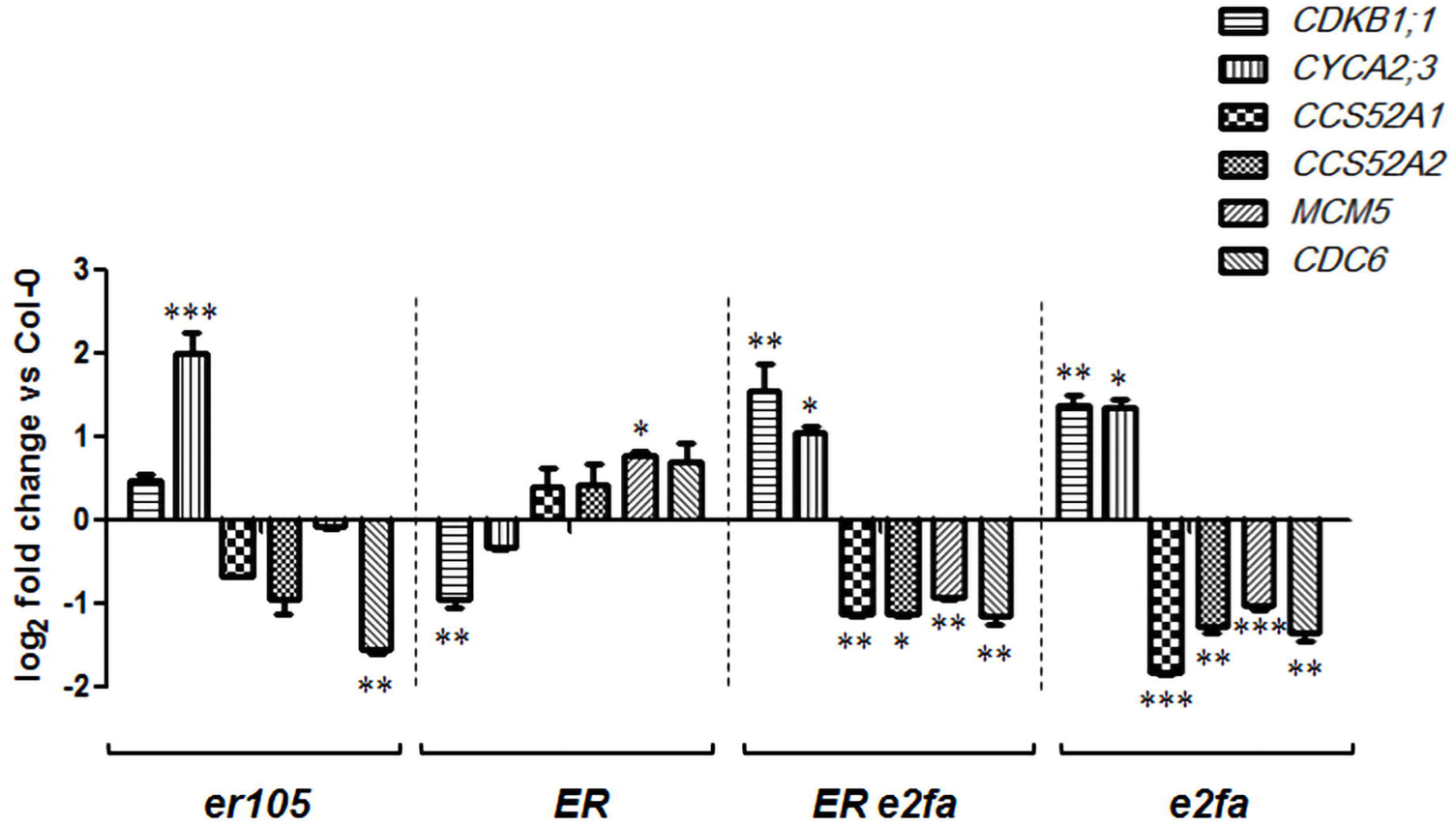
Expression level of E2Fa target genes are changed in *ERECTA* mutants. Values were normalized to *Arabidopsis Ubiquitin5* expression levels and represented as n - fold compared to the wild type (Col-0). All assays were carried out at least three times and levels of statistical significance were calculated using student’s *t*-test (* P≤ 0.05, **P ≤ 0.01, ***P ≤ 0.001).

### ERECTA interacts with E2Fa via its kinase domain

To investigate how ERECTA regulates E2Fa, we tested whether they physically interact with each other by using *in vitro* and *in vivo* interaction assays. The interaction between ERECTA kinase domain (ERK) and E2Fa was verified in pulldown assay (Fig. 6a) and yeast-two-hybrid assay (Fig. 6b). The co-immunoprecipitation (Co-IP) experiment demonstrated that the intact ERECTA interacts with E2Fa *in vivo* (Fig. 6c).

**Figure 6.**
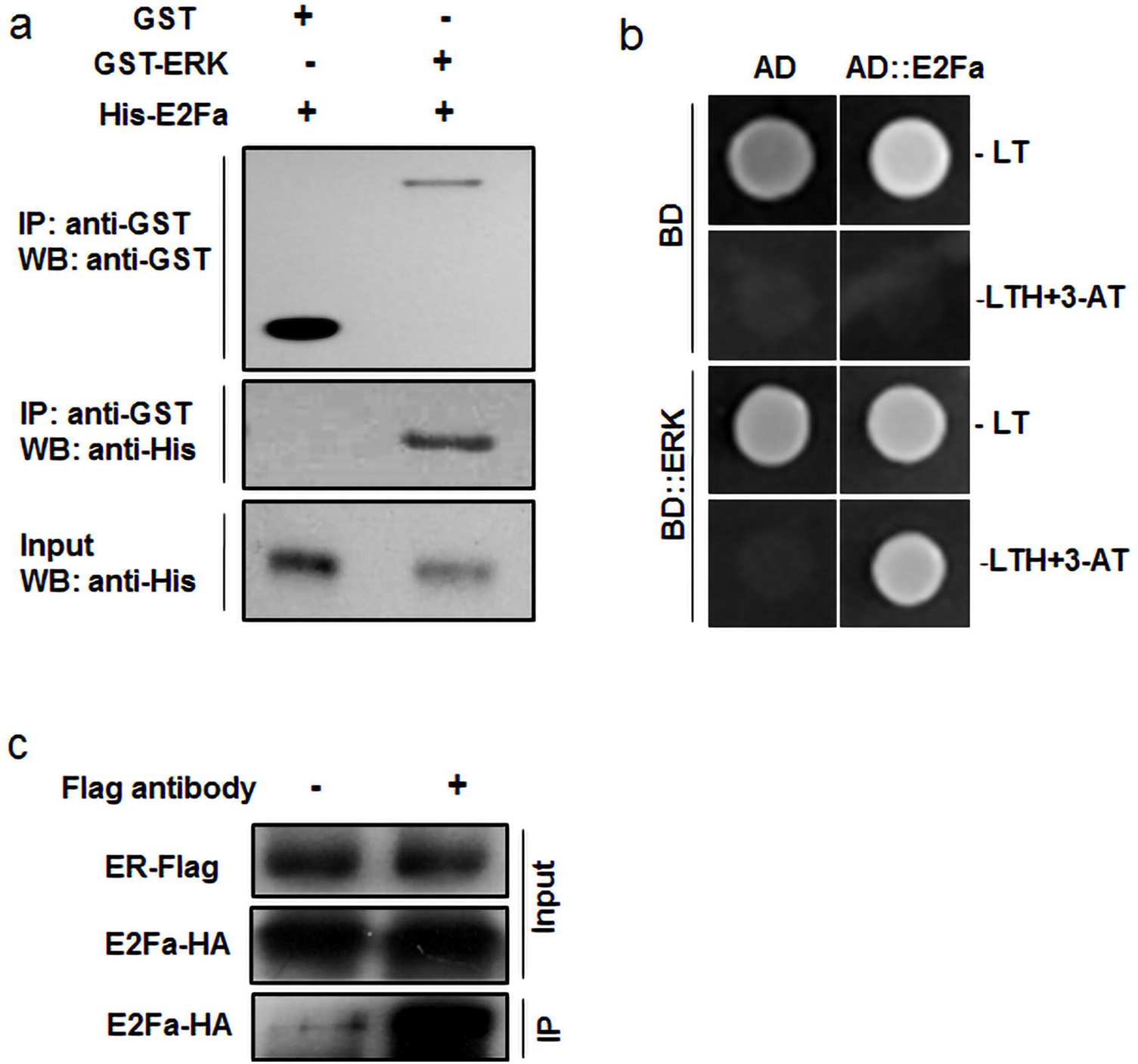
ERECTA interacts with E2Fa via its kinase domain (ERK). (a)ERECTA kinase domain binds E2Fa *in vitro* by pull-down assay. ERECTA kinase domain (amino acids 615 – 975) fused with GST tag, E2Fa fused with 6×His-tag, were expressed and purified from *E. coli.* His-E2Fa was incubated with GST or GST-ERK and pulled down with glutathione agarose beads. Bound proteins were detected by immunoblotting with anti-His antibody and anti-GST antibody. (b)ERECTA kinase domain can bind E2Fa *in vivo*. In yeast-two-hybrid assay, interaction was indicated by the ability of cells to grow on synthetic dropout medium lacking Trp/Leu/His and containing 10 mM3-aminotriazole. The Gal4 DNA binding domain was fused with ERECTA kinase domain (shown as BD⸬ERK) and the Gal4 activation domain was fused with E2Fa (shown as AD⸬E2Fa). The Gal4 DNA binding domain expressed by pGBKT7 (shown as BD-) and the Gal4 DNA activation domain expressed by pGADT7 (shown as AD-) were used as negative controls. (c)The interaction between ERECTA and E2Fa was detected by co-immunoprecipitation assay. *Arabidopsis* mesophyll protoplasts were transfected with ERECTA-Flag together with E2Fa-HA, and the extracted proteins were immunoprecipitated by anti-Flag antibody. Gel blots were probed with anti-HA antibody. IP, immunoprecipitation; WB, western blot.

## Discussion

Plant WUE is critical to plant survival and crop yield. In this study, we provided evidence to suggest an EDT1/HDG11-ERECTA-E2Fa genetic pathway that regulates plant WUE. The homeodomain transcriptional factor EDT1/HDG11 upregulates *ERECTA*, ERECTA in turn interacts with E2Fa to promote endoreduplication, leading to higher ploidy level and thus larger cell and reduced stomatal density, ultimately improved WUE. *AtEDT1/HDG*11-conferred drought resistance and improved WUE have been demonstrated by overexpressing *AtEDT1/HDG11* in many plant species including monocots and woody plant (Yu et al., 2008; Cao et al., 2009; Ruan et al., 2012; Yu et al., 2013; Zhu et al., 2015; Li et al., 2016; Yu et al., 2016; Zhu et al., 2016; Liu et al., 2017), suggesting that the underlying mechanism might be well conserved in higher plants.

*Arabidopsis ERECTA* and its functional homologs, *ERL1* and *ERL2*, show synergistic interaction in promoting above ground organ growth (van Zanten et al., 2009). However, *ERECTA, ERL1*, and *ERL2* have subtle different roles in epidermal development (Shpak et al., 2005; Bergmann and Sack, 2007; Lee et al., 2012; Pillitteri and Torii, 2012). Elevated expression of *ERECTA* leads to an increased epidermal cell size and reduced stomatal density in leaves, but the stomatal index is not significantly changed, consistent with the report that *er* single mutation displayed an increased stomatal density and smaller epidermal cells, without significant change in stomatal index (Masle et al., 2005; Shen et al., 2015). Detailed analysis of higher order mutant combinations revealed that ERECTA primarily acts at the early steps of stomatal development to suppress entry asymmetric divisions (Pillitteri and Torii, 2012).

Our work revealed that the activated transcription of *ERECTA* by EDT1/HDG11 modulates stomatal density mostly through epidermal pavement cell growth rather than in stomatal development. Cell growth is associated with the increase of ploidy levels (Figure 1 and 2). Considering that the phenotype of overexpressed *EDT1/HDG11* is largely blocked by knocking out of *ERECTA* in double mutant *edt1er-105*, we concluded that *ERECTA* is a major target of EDT1/EDT1/HDG11 for augmenting leaf cell size and decreasing stomatal density, thereby improving plant WUE in *edt1*.

Moreover, the negative correlation between stomatal density and WUE is nicely demonstrated by using Arabidopsis *epf* mutants with altered stomatal density or by genetic manipulation of *EPF* genes (Doheny-Adams et al., 2012; Franks et al., 2015; Hepworth et al., 2015). Reduced stomatal density improves drought tolerance without yield penalty in barley (Hughes et al., 2017).

Taken together, these results suggest that leaf cell size is positively correlated with WUE. Enlarged leaf cells might have higher metabolic activities including photosynthesis rate contributing to improved WUE, which is in line with Kleiber’s law (Kleiber, 1932)

Previous studies have demonstrated key functions of *ERECTA* family genes in regulating the architecture of whole plant and the development of stomata (van Zanten et al., 2009; Pillitteri and Torii, 2012). Here we report a role of *ERECTA* for cell size and stomatal density regulation, which coordinates with other developmental processes to optimize water use efficiency. Notably, *ERECTA* family genes regulate the architecture of whole plant and the development of stomata mainly through the MAPK signaling cascade, influence the stomatal development and patterning by phosphorylating and regulating the stability of transcription factors SPCH, MUTE and FAMA (Lampard et al., 2008; Lampard et al., 2009; Meng et al., 2012). These development regulations influence cell fate determination and have drastic effects on the growth, sometimes even cause death or sterility of plants. However, *ERECTA* gene increases cell size and decreases stomatal density through transcription factor E2Fa, influencing cell cycle and endocycle transition, which modifies the stomatal density without affecting the cell fate determination.

E2F transcription factors are required for the progression in both mitotic and endocycle (De Veylder et al., 2002). E2Fa is localized to both the cytoplasm and nucleus (Kosugi and Ohashi, 2002; Magyar et al., 2012), while ERECTA is localized in the plasma membrane (Horst et al., 2015). Although we demonstrated that E2Fa is a downstream target of ERECTA (Fig. 4 and 5), how ERECTA transduces the signal to E2Fa awaits further investigation. As a receptor-like protein kinase, ERECTA likely relays the signaling through phosphorylation of its targets. We attempted to demonstrate whether ERECTA directly phosphorylates E2Fa. ERECTA and E2Fa can physically interact *in vitro* and *in vivo* (Fig. 6), but immunoprecipitated endogenous ERECTA from *Arabidopsis* plants failed to directly phosphorylate E2Fa *in vitro* (data not shown), suggesting that the interaction between ERECTA and E2Fa may cause modifications of E2Fa other than phosphorylation, or its phosphorylation depends on other protein factors and may require signals of development or/and environmental cues. Further research is needed to identify these components.

In conclusion, we have demonstrated that EDT1/HDG11 transcriptionally activates *ERECTA*. ERECTA interacts with E2Fa to modify E2Fa activity, which regulates the expression of E2Fa target genes in the mitosis-to-endocycle transition and allows cells to decrease their division rate and promote endoreduplication. Consequently, these changes result in increased leaf cell size, reduced stomatal density, and improved water use efficiency.

## Methods

### Plant materials and growth conditions

*Arabidopsis thaliana* Columbia-0 ecotype (Col-0) was used as wild type. All mutants and transgenic plants used in the present work are in the Col-0 background. Some plant materials used in this study were previously described: *edt1* (Yu et al., 2008), *er-105*(Shpak et al., 2005), *35S⸬HA-HDG11* (Xu et al., 2014), *e2fa*, *HA-E2Fa* (Xiong et al., 2013). CS89504 (*er-105*) and CS855653 (*e2fa*) mutant seeds were obtained from Arabidopsis Biological Resource Center. Double mutants were generated by genetic crosses. Plants of a correct genotype were isolated from the F2 populations. The activated expression of *EDT1/HDG11* in *edt1*, and overexpression of *ERECTA* in *ER* lines, conferred resistance to Basta. Thus, progenies of each cross were first tested for Basta resistance, and subsequently the genotype of individual plants, homozygous lines were identified by PCR-based genotyping, the presence of the *er-105* mutation was determined by RT-PCR using the primer pairs: ERECTA-RT-F and ERECTA-RT-R (see Supplemental Table 1). *Arabidopsis* seeds were surface sterilized for 10 □min in 10% bleach and washed at least five times with sterile water. Plant seeds were kept at 4 □°C for 2 days in darkness before germination on horizontal agar plates containing solid Murashige and Skoog (MS) medium with 1% (w/v) sucrose at 22 □°C under 16-h light/8-h dark cycles. Adult plants grow in pit moss under long-day conditions (16-h light / 8-h dark) at 22–24°C.

### DNA constructs and plant transformation

To prepare the ERECTA overexpression construct, the primer set gERECTA-F and gERECTA-R (Supplemental Table 1) and Full-length genomic coding regions for ERECTA fragments were amplified and cloned into pD0NR207, and subsequently shuttled it into the expression binary vector pCB2004 (Lei, 2007). To make the 35Spro:ERECTA-Flag construct, which was used for Co-IP assays. The full-length ERECTA coding region was amplified by PCR from total cDNA, fused to the FLAG tag at the 3’ end of the gene, and cloned between a 35S-driven promoter and NOS terminator (Xiong and Sheen, 2012). All of these constructs utilized the gateway system technology. These constructs were then individually transformed into *Agrobacterium tumefaciens* strain (C58C1), and introduced into *Arabidopsis* plants by the floral dip method.

### Cytological analysis, WUE analysis, and Flow cytometry analysis

To measure cell density and stomatal density, single optical sections of the adaxial side of PI-stained cotyledons from the wild type and the mutants were acquired with a 40×objective. For statistical analysis of cell density and stomatal density, 30 leaves were sampled per genotype.

Photosynthesis (P) and transpiration (T) rates were measured on fully developed leaves of plants grown in a greenhouse, using a Li-6400 portable gas-exchange system (LI-COR, Lincoln, NE, USA). All measurements were conducted between 9:00 a.m. and 11:00 a.m. Conditions in the LI-6400XT chamber were as follows: constant air flow rate, 50μmol/s; CO_2_ concentration, 400 μmol/mol; temperature, 23± 2 °C; relative humidity (RH), 75-80%; and photosynthetic photon flux density, 50μmol (photon)/m^2^ per s. Incoming air was aspirated through a custom-made humidifier consisting of a plastic bottle with wet paper towels. Gas exchange measurements were taken after steady state had been reached. WUE was defined as P/T ratio and derived from the measured P and T.

For flow cytometry measurements, the fifth rosette leaf was collected and chopped with razor blades in nuclei extraction buffer and stained with propidium iodide (PI) as described before (Galbraith et al., 1983; Jing et al., 2009). The experiment was performed three times independently. A total of 10,000 nuclei were measured per analysis. Flow cytometry data were obtained using a FACS calibur flow cytometer (BD Biosciences).

### qRT-PCR gene expression analysis

For verification of microarray results, the RNA were isolated using Trizol reagent (Invitrogen) from four-weeks-old rosette leaves of wild type and *edt1* used for the microarray experiment. For gene expression analysis, RNA was prepared from whole seedlings (7 days after germination) using Trizol reagent (Invitrogen). RNA was reverse transcribed to cDNA using PrimeScript™ RT reagent Kit (Perfect RealTime) (TAKARA), according to the manufacture’s procedure. An aliquot of μL of the synthesized cDNA was used as template in a 10μL PCR reaction with gene specific primers (Supplemental Table 1) and the SYBR^®^ Premix Ex Taq™ II (TliRNaseH Plus) kit (TAKARA). *Ubiquitin5*(*UBQ5*, At3g62250) was used as the internal control. Real time PCR is running on Stepone real-time PCR systems (Applied Biosystems).

### Confocal microscopy analysis

To visualize epidermal cell outlines, seedlings were stained with 0.2 mg/ml propidium iodide for 20 min and washed twice in water. The fluorescence of Propidium iodide fluorescence was imaged under a fluorescence microscope (ZEISS Axioskop 2 plus): 543-nm of the laser was used for excitation, and emission was detected at 620 nm.

### Yeast-one-hybrid and Yeast-two-hybrid analysis

Yeast strain YH187 and destination vectors (pHIS2 and pGADT7); Yeast strain YHA109 and destination vectors (pGADT7 and pGBKT7) were obtained from Jin- Song Zhang (Chinese Academy of Sciences). For the yeast-one-hybrid analysis, the prey vector harbouring *HDG11* (pAD-HDG11) and the bait vector pHIS2 harbouring the *ERECTA* promoter fragment *cis*-element 1 or *cis*-element 2 were used to transform yeast cells. The transformants were observed for their growth on SD/-Leu-His medium or this medium plus 3-aminotriazole (3-AT) following the instruction of the BD Matchmaker Library Construction & Screening Kits (www.bdbiosciences.com). Transformation with empty vectors pGADT7 and pHIS2 was used as negative controls. The experiments were repeated three times with the same results. For the yeast-two-hybrid analysis, the kinase domain of ERECTA was inserted into pGBKT7 vector to construct the bait plasmid, while the prey plasmid was constructed with pGADT7 vector and the full-length CDS of E2Fa. Positive clones were screened using SD/-Leu-Trp-His with 3-AT. The primers could be found in Supplemental Table 1.

### Pull-down assay

To generate recombinant proteins, cDNA fragments of E2Fa was cloned into the pET15b vector (Invitrogen), The kinase domain of ERECTA was amplified and ligated into the pGEX-5X-1 vector. His-E2Fa and GST-ERK (ERECTA kinase domain) fusion protein were expressed using bacterial expression system and purified. GST-ERK fusion protein was incubated with Glutathione-Superflow Resin (Clontech) at 4°C for 2 h, and the GST tag was used as a negative control. The beads were cleaned with washing buffer for four times. Then the beads were incubated with His-E2Fa at 4°C for 2 h respectively. The beads were cleaned with washing buffer for four times. Western blot was used to detect the SDS-PAGE separation results of pulled-down mixtures in nitrocellulose membrane with anti-His antibody and anti-GST antibody (Abmart).

### Co-immunoprecipitation (Co-IP) for the E2Fa-ERECTA interaction assay

Co-IP assays were performed as described (Xiong et al., 2013). Protoplasts (5 × 10^5^) were transformed with 15μg ER-Flag and 15μg E2Fa-HA, then incubated in 5 ml of mannitol (0.5 M) and KCl (20 mM) buffer (4 mM MES, pH 5.7) in Petri dishes (100 mm × 20 mm, 10^5^ cells/each) for 4 hours. Collected protoplasts were lysed in 500μl Co-IP buffer (400 mM HEPES pH 7.4, 2 mM EDTA, 10 mM pyrophosphate, 10 mM glycerol phosphate, 0.3% CHAPS and 1X cocktail inhibitors [Roche]). To immunoprecipitate ERECTA, protein extracts were incubated without/with anti-Flag antibody at 4°C for 2 h, and additional 2 h after adding 15 μl protein G sepharose beads (GE healthcare). The immunoprecipitated proteins were washed four times with low salt wash buffer (400 mM HEPES pH 7.4, 150 mM NaCl, 2 mM EDTA, 10 mM pyrophosphate, 10 mM glycerol phosphate, 0.3% CHAPS) before SDS-PAGE separation and protein blot analyses.

### ChIP-qPCR assay

The ChIP experiment was performed as described (Mukhopadhyay et al., 2008) with minor modifications. One gram of leaves of four-weeks-old wild type and the transgenic plants harboring *35Spro⸬HA-HDG11* (Xu et al., 2014) were harvested and immersed in 1% formaldehyde under vacuum for 10 min. Glycine was added to a final concentration of 0.125 M, and incubation was continued for 5 min. After washing, the seedlings were ground into a fine powder with liquid nitrogen and resuspended in nuclei isolation buffer. The nucleus were then collected by centrifugation and resuspended with nuclei lysis buffer. The crosslinked DNA/protein complexes were fragmented by sonication with an Ultrasonic Process (Sonics), resulting in fragments of ≤ 500 bp. After centrifugation (16,000g), the supernatant, 30μL was used as input, while the rest was divided into two samples, of which one sample was treated with an HA tag-specific monoclonal antibody (1:100 for ChIP assay, HA-Tag, 26D11, Mouse mAb, M20003, Abmart, Shanghai, China) and the other with no antibody. The samples were incubated overnight. Immunoprecipitates were collected with 10μL of supernatant was precleared with 50μL of the protein A agarose/salmon sperm DNA (Millipore) and subsequently eluted from the beads. All centrifugation steps with bead-containing samples were done at 1000g. Proteins were de-cross-linked, and DNA was purified by phenol/chloroform/isoamyl alcohol extraction and ethanol precipitation. Pellets were resuspended in 40μL of Tris-EDTA buffer (0.05 M Tris-HCl and 0.02 M EDTA, pH 6.5). Input and immunoprecipitated chromatin were used for qRT-PCR analysis using various *ERECTA* promoter-specific primers that were designed to amplify *ERECTA* promoter fragments (see Supplemental Table 1). The qRT-PCR analysis was performed according to the procedure described previously (Mukhopadhyay et al., 2008). The regions of ERECTA 3’ UTR that do not contain putative binding sites of HD class transcription factors (*cis* NC) was as negative control. The relative quantity value is presented as the DNA binding ratio (differential site occupancy).

### Accession Numbers

*EDT1/HDG11* AT1G73360, *ERECTA* AT2G26330, *E2Fa* AT2G36010, *ERL1* AT5G62230, *ERL2* AT5G07180.

## Author Contributions

C.X., Y.X., and X.G. designed the experiments. X.G., Y.W., P.X., G.Y., Y.X., and L.Z. performed the experiments and data analyses. X.G. wrote the manuscript. C.X supervised the project and revised the manuscript.

## Acknowledgements

This study was supported by grants from the National Nature Science Foundation of China (30830075), the Ministry of Science and Technology of China (“973201D Program 2012CB114304), the Chinese Academy of Sciences (KSCX3-YW-N-007). We thank ABRC for providing the mutant seeds.

## Supplemental information

Supplemental Figure 1. Growth phenotypes and photosynthesis of wild type, *edt1*, *er-105* and *edt1er-105*.

Supplemental Figure 2. *ERECTA* but not *ERL1* or *ERL2* is transcriptionally upregulated by EDT1/HDG11.

Supplemental Figure 3. Growth phenotypes of wild type, *ER*, *e2fa* and *ERe2fa*.

Supplemental Table 1. Primers used in this study.

